# Centromere-specific antibody-mediated karyotyping of Okinawan *Oikopleura dioica* suggests the presence of three chromosomes

**DOI:** 10.1101/2020.06.23.166173

**Authors:** Andrew W. Liu, Yongkai Tan, Aki Masunaga, Charles Plessy, Nicholas M. Luscombe

**Author notes:** Equal contribution (Andrew W. Liu, Yongkai Tan). **<u>Corresponding authors:</u>**.

## Abstract

*Oikopleura dioica* is a ubiquitous marine tunicate of biological interest due to features that include dioecious reproduction, short life cycle, and vertebrate-like dorsal notochord while possessing a relatively compact genome. The use of tunicates as model organisms, particularly with these characteristics, offers the advantage of facilitating studies in evolutionary development and furthering understanding of enduring attributes found in the more complex vertebrates. At present, we are undertaking an initiative to sequence the genomes of *Oikopleura* individuals in populations found among the seas surrounding the Ryukyu Islands in southern Japan. To facilitate and validate genome assemblies, karyotyping was employed to count individual animals’ chromosomes *in situ* using centromere-specific antibodies directed against H3S28P, a prophase-metaphase cell cycle-specific marker of histone H3. New imaging data of embryos and oocytes stained with two different antibodies were obtained; interpretation of these data lead us to conclude that the Okinawan *Oikopleura dioica* has three pairs of chromosomes, akin to previous results from genomic assemblies in Atlantic populations. The imaging data have been deposited to the open-access EBI BioImage Archive for reuse while additionally providing representative images of two commercially available anti-H3S28P antibodies’ staining properties for use in epifluorescent and confocal based fluorescent microscopy.

## Introduction

Karyotyping is a long-established histochemical method to visualize chromosomes of eukaryotes (Tjio & Levan, 1950; Hsu & Benirschke, 1967). A multi-dye reagent developed at the turn of the 20^th^ century for the diagnosis of infections in human histological preparations (Giemsa 1902; 1904) was later used to stain chromosomes themselves in order to study their numbers, translocations, and other aberrations. This rapid technique, involving the use of stains including methylene blue, eosin, and azure B allows for observation of chromosomes with a simple light microscope, naturally lending itself to a first attempt for karyotyping analysis.

Although individual chromosomes have been resolved by histochemical techniques in *O. dioica*, the reported results differ in numbers from n=3 (Körner, 1952) to n=8 (Colombera & Fernaux, 1973). More recently, metaphase-specific histone 3 (H3) markers have been used to determine the structure and the segregation of genetic material during oogenesis *in situ* (Ganot *et al.*, 2006; Schulmeister *et al.*, 2007) while providing greater detail and resolution. One such marker is histone H3 phosphorylated at Ser-28 (Kawajiri *et al.*, 2003); although it is typically used to identify centromeres during metaphase (Kurihara *et al.*, 2006), we observed in data presented in previous studies that signals were not confined to centromeres. More importantly, the localization of the H3S28P signal depends on the phase of the cell cycle: spatially punctate signals were found evenly spread within the nuclear envelope during prophase, while condensed chromatin gave an outlined staining of the sister chromatids during metaphase in a manner consistent with alignment along the metaphase plate (Table 1; Campsteijn *et al.*,, 2012; Olsen *et al.*,, 2018; Feng & Thompson, 2018; Feng *et al.*,, 2019). Moreover, a structure in which genetic material is sequestered in a ∏-shaped conformation has been observed during meiotic cell divisions between the final phases of oogenesis and mature oocytes (Ganot *et al.* 2008). However, these results were all obtained from the same laboratory strain originating from the Atlantic Ocean. Considering the discrepancy of past findings, and the fact that our laboratory strain originates from a geographically distinct ocean, we applied H3S28P staining on intact embryos and oocytes to confirm the chromosome count and validate our genome sequencing assemblies of Okinawan *O. dioica* marine populations among the Ryukyu Islands of southern Japan.

**Table 1.**
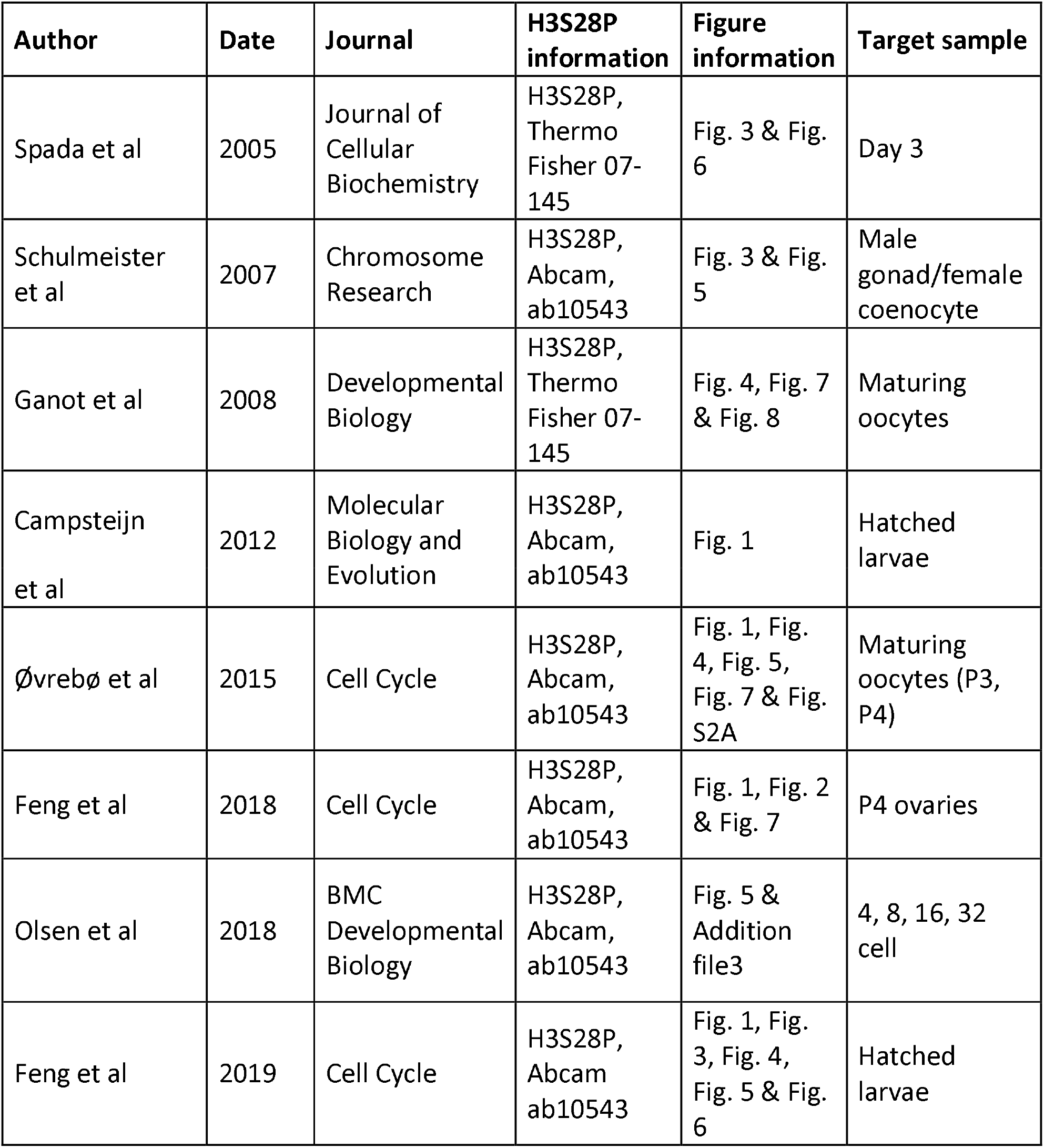
Reference to images cited in this study.

| **Author** | **Date** | **Journal** | **H3S28P information** | **Figure information** | **Target sample** |
| --- | --- | --- | --- | --- | --- |
| Spada et al | 2005 | Journal of Cellular Biochemistry | H3S28P, Thermo Fisher 07-145 | Fig. 3 & Fig. 6 | Day 3 |
| Schulmeister et al | 2007 | Chromosome Research | H3S28P, Abcam, ab10543 | Fig. 3 & Fig. 5 | Male gonad/female coenocyte |
| Ganot et al | 2008 | Developmental Biology | H3S28P, Thermo Fisher 07-145 | Fig. 4, Fig. 7 & Fig. 8 | Maturing oocytes |
| Campsteijn et al | 2012 | Molecular Biology and Evolution | H3S28P, Abcam, ab10543 | Fig. 1 | Hatched larvae |
| Øvrebø et al | 2015 | Cell Cycle | H3S28P, Abcam, ab10543 | Fig. 1, Fig. 4, Fig. 5, Fig. 7 & Fig. S2A | Maturing oocytes (P3, P4) |
| Feng et al | 2018 | Cell Cycle | H3S28P, Abcam, ab10543 | Fig. 1, Fig. 2 & Fig. 7 | P4 ovaries |
| Olsen et al | 2018 | BMC Developmental Biology | H3S28P, Abcam, ab10543 | Fig. 5 & Addition file3 | 4, 8, 16, 32 cell |
| Feng et al | 2019 | Cell Cycle | H3S28P, Abcam ab10543 | Fig. 1, Fig. 3, Fig. 4, Fig. 5 & Fig. 6 | Hatched larvae |

## Methods

### *Oikopleura dioica* culture, staging & preparation of biological material

#### Histochemical staining

Live specimens were collected from Ishikawa Harbor (26 °25’39.3 “N, 127 °49’56.6 “E) by a hand-held plankton net and cultured in the lab (Masunaga *et al.* 2020). Mature females were collected prior to spawning, individually washed with filtered autoclaved seawater (FASW) 3 times for 10 minutes and placed in separate 1.5 ml tubes containing 500 μl of FASW. Nearly mature males, full of sperm, were also washed 3 times in FASW. Mature males that successfully made it through the washes intact were placed in 100 μl of fresh FASW and allowed to spawn naturally. As soon as females spawned, each individual clutch of 100-200 eggs was washed three times for 10 minutes by moving eggs along with a pulled capillary micropipette from well to well in a 6-well dish, each containing 5 ml of FASW, and left in a fresh well of 5 ml FASW in the same dish. These were stored at 17 °C and set aside for fertilization. Staged embryos were initiated by gently mixing 10 μl of the spawned male sperm with the awaiting eggs in FASW at 23 °C. Developing embryos were staged and collected by observation under a dissecting microscope. These embryos were quickly dechorionated using 0.1% sodium thioglycolate and 0.01% actinase in FASW for 2-3 minutes, then promptly washed with 2 washes with filtered autoclaved seawater prior to fixation and staining. Unfertilized eggs were treated similarly.

Embryos were Giemsa stained as previously described in Shoguchi *et al.*, 2005.

#### Immunostaining

Washed eggs and embryos were immediately fixed in 4% w/v paraformaldehyde, 100 mM MOPS pH 7.5, 0.5 M NaCl, 0.1% triton-X100 at 23 °C ON (Campsteijn et al, 2012). The samples were then washed for 10 minutes once with PBSTE (PBS supplemented with 1 mM EDTA) and 3 times for 10 min with PBSTEG (PBS supplemented with 1 mM EDTA and 0.1 M glycine). The samples were blocked using PBSTE supplemented with 3% bovine serum albumin at 4 °C overnight. Rabbit polyclonal (Thermo Fisher Scientific Cat# 720099, RRID:AB_2532807) or rat monoclonal (Abcam Cat# ab10543, RRID:AB_2295065) primaries directed against H3S28P were diluted 1:100 in PBSTE 3% BSA and incubated at 4 °C for 3 days. The next morning, these were washed in PBSTE for 10 minutes 3 times and incubated with anti-rabbit (Thermo Fisher Scientific Cat# A-11034, RRID:AB_2576217) or anti-rat (Molecular Probes Cat# A-11006, RRID:AB_141373) Alexa488 conjugated secondary antibodies diluted 1:500 with PBSTE 3% BSA at 4 °C ON. The following morning, samples were washed 3 times for 10 min with PBSTE. The samples were mounted on cleaned glass slides (Matsunami Glass, S2441) with fluorescence preserving mounting medium (ProLong. Fluoromount G Mounting Medium, RRID:SCR_015961) covered with No.1 35 × 50 mm glass coverslips (Matsunami Glass, C035551) and sealed with nail polish.

#### Image acquisition

Both a Nikon Ni-E epifluorescent and a Zeiss LSM 510 Meta confocal microscopes were used to acquire Z-stack images of eggs and embryos. Brightfield images were obtained using a 20x/0.75 CFI Plan Apo λ objective (Nikon, MRD00205) for histochemical staining. Epifluorescent immunofluorescent images were obtained with both 20x/0.75 and 40x/0.95 CFI Plan Apo λ air objectives (Nikon, MRD00405); each sample acquisition was Z-stacked with each plane set at an interval of 1 μm. Confocal images were acquired using a 40x/0.75 EC Plan-Neofluar M27 (Zeiss, 420360-9900-000) and 63x/1.4 Plan-Apochromat M27 oil immersion (Zeiss, 420782-9900-79) objectives; each sample acquisition was Z-stacked, line averaged twice with each plane set at an interval of 0.6 and 0.27 μm, respectively.

#### Image processing and analysis

Images acquired from a Nikon Ni-E epifluorescent were deconvoluted with Nikon Elements software. Images for both epifluorescent and confocal acquisitions were analyzed using Imaris software `SPOT DETECTION` tool (Imaris, RRID:SCR_007370) for embryos and unfertilized eggs, parameters set at 0.5 and 0.43 μm spot detection size, respectively, and software preset to `QUALITY` auto signal threshold for each individual cell within a sample. Epifluorescent and confocal acquisitions of embryos and their subsequent analysis were performed independently by different researchers to exclude bias.

## Results

Initial attempts at visualizing individual chromosomes were done with developing embryos and Giemsa staining. The spreads from two time points, 32- and 64-cell developmental stages, gave results with counts ranging between 11-27 stains per cell (BioImage Archive, S-BIAD21, Experiment A). Although hypotonic-induced cell spreads were confined as a result of incomplete dechorionation and digestion with the enzymatic dissociation cocktail, groups of chromosomes were easily associated to a single cell. However, individual chromosomes were difficult to resolve due to the low resolution of images. In order to eliminate possible miscounts and other Giemsa staining artifacts, immunostaining was used to count individual chromosomes using a centromere-specific primary antibody directed against H3S28P and a secondary antibody conjugated to Alexa488 directed against the primary antibody.

Signal-based thresholding was employed to determine the number of distinct 515 nm emission signals present in acquired images from epifluorescent and laser confocal microscopes (BioImage Archive, S-BIAD21, Experiment B & D). The data was analyzed using the Imaris `SPOT DETECTION` tool (Oxford Instruments). Two types of nuclei were apparent within each embryo: nuclei containing evenly distributed, clearly separated spots that were interpreted as being in prophase (Figure 1A and 1B, blue circles) and nuclei with intense clusters of signals in the center, considered to be in metaphase (Figure 1A and 1B, red squares). Counts from these two classes of nuclei fall into separate distributions (Figure 1C and 1D). Both epifluorescent and confocal acquisitions were in near agreement, epifluorescence n = 20, mean 6.2, 95% CI 5.6 - 6.8; confocal n = 13, mean 6.4, 95% CI 5.7 - 7.1 and epifluorescence n = 20, mean 12, 95% CI 11.0 - 13.0; confocal, n = 14, mean 14.1, 95% CI 12.9 - 15.3. We interpret the results as a count of 12 distinct centromeres in prophase cells and a count of 6 larger spots identifying pairs of centromeres in metaphase (Figure 1B).

**Figure 1:**
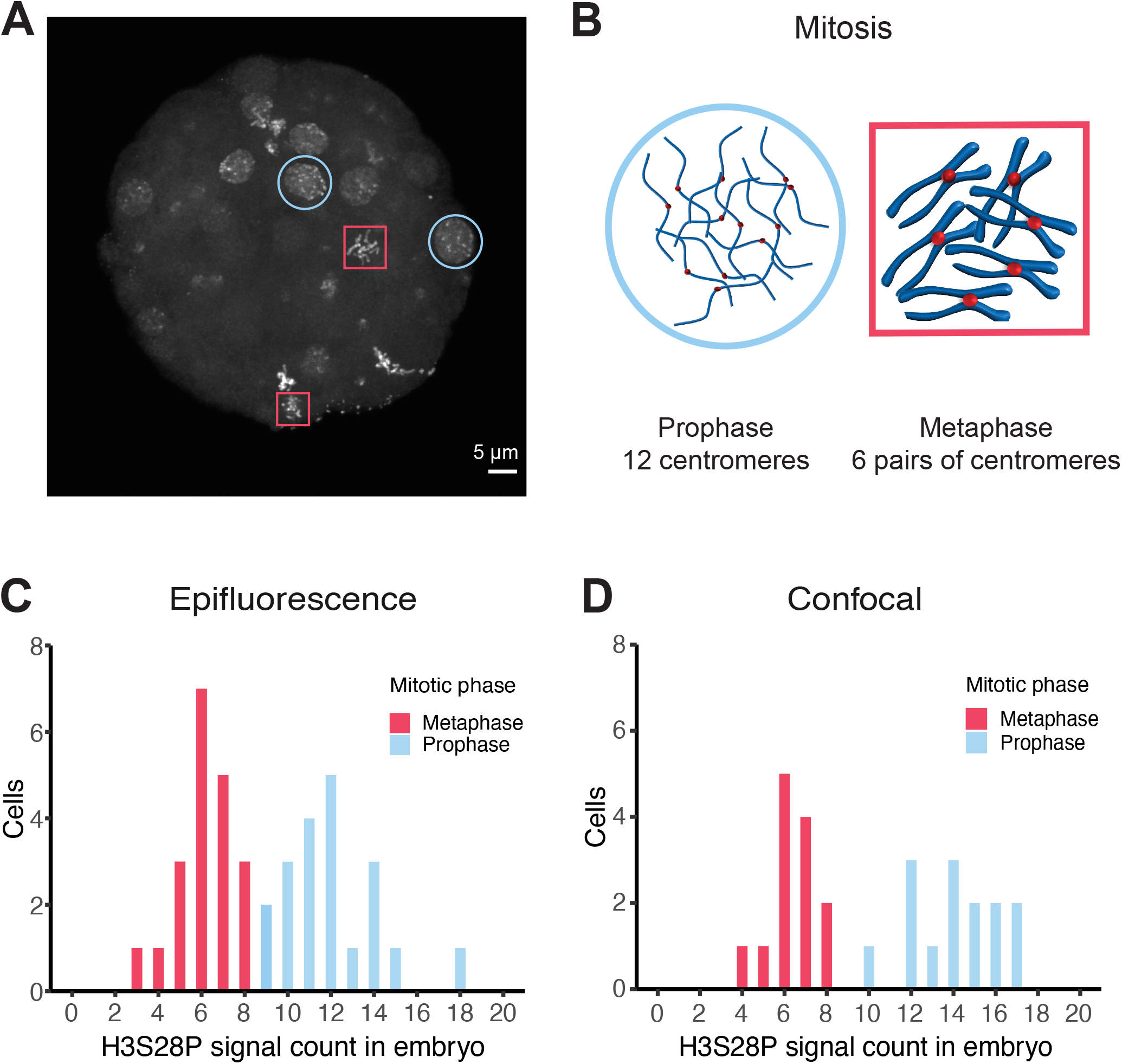
Centromere counts from embryos. Anti-H3S28P rabbit-derived polyclonal stained 64-cell whole-embryo chromosomal imaging data analyzed by Imaris software `SPOT DETECTION` tool using different microscopy techniques. **A** Maximum projection of confocal image of an embryo demonstrating the differences in signal localization appearance and signal count, which was inferred to represent distinct cell cycle phases. Red box, metaphase; blue circle, prophase. **B** Schematic interpretation of signals with respect to chromatin structure during prophase and metaphase cell cycle states. As a simplification, all chromosomes have been drawn at an equal length although they actually vary in *O. dioica*. **C** Distribution of signal counts within individual cells using epifluorescent (n = 40) and **D** confocal (n = 27) microscopes. Two distinct populations were observed in a bimodal distribution, which corresponded with cell cycle stage. Red, metaphase; blue, prophase.

To confirm our observations on germ cells and therefore rule out polyploidy, which is frequent in *O. dioica’s* somatic cells (Ganot & Thompson, 2002), we also analyzed oocytes in prometaphase I before fertilization (Schulmeister *et al.*, 2007). We identified confined groupings of signals in unfertilized eggs (Figure 2A; BioImage Archive, S-BIAD21, Experiment E). Images were analyzed using the Imaris `SPOT DETECTION` tool to determine chromosome counts and their distributions (Figure 2B). Counts from the compact rosette-shaped genetic material averaged near 6 (n = 23, mean 5.70, 95% CI 5.2 - 6.2). Visual inspection of individual Z-sections (Figure 2C) confirm agreement with the Imaris count analysis and annotation (Figure 2D). We interpret these results as each spot corresponding to a pair of centromeres from paired chromatids forming a synapsis in unfertilized eggs (Figure 2E).

**Figure 2:**
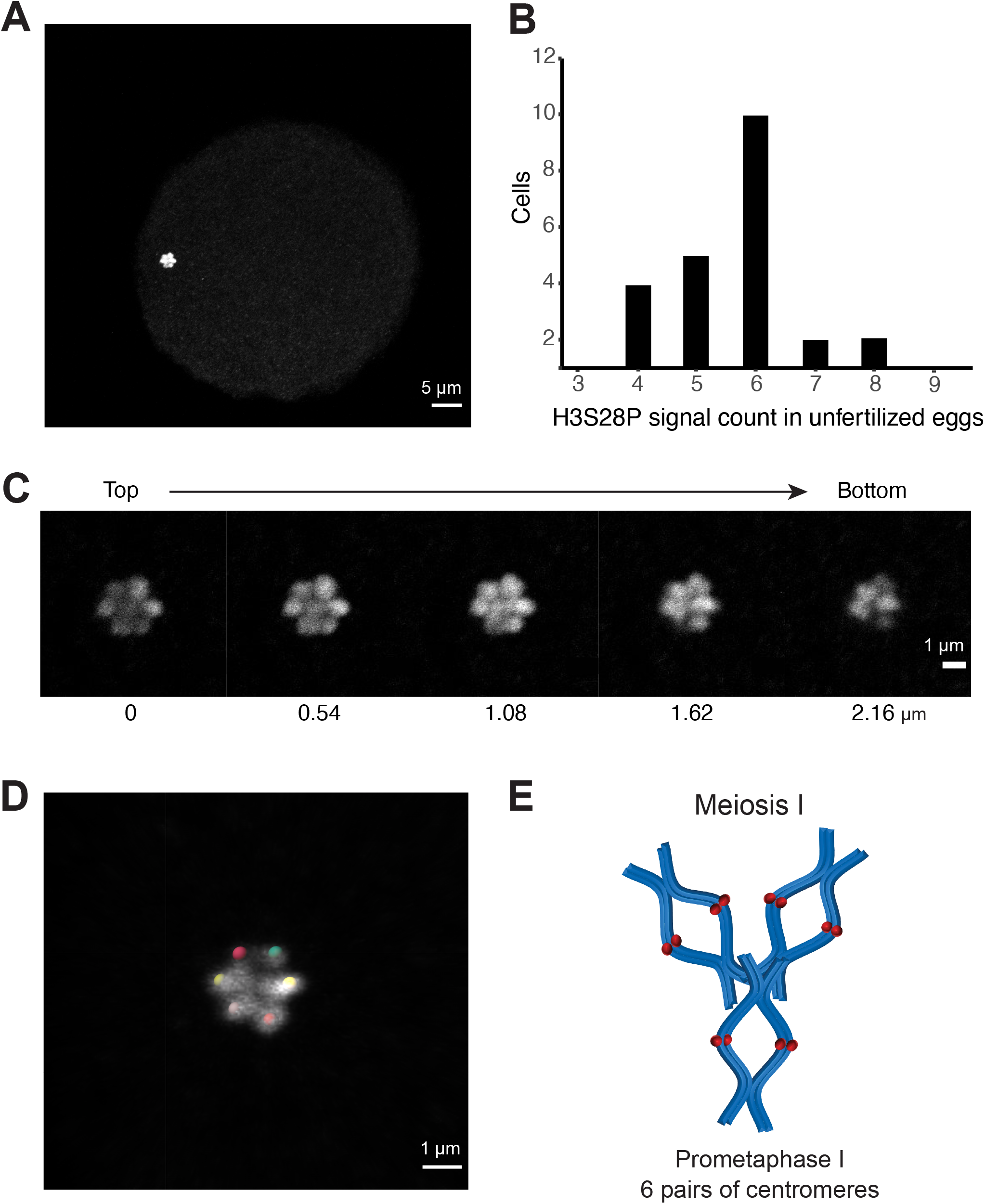
Centromere counts from unfertilized eggs. **A** Maximum signal projection of a representative confocal Z-stack acquisition of anti-H3S28P rat monoclonal stained oocyte used for the count analysis. **B** Distribution of signal counts from centromere-stained oocyte genetic material, analyzed by Imaris software `SPOT DETECTION` tool (n = 23). **C** Individual Z-sections from same image acquisition showing the 3D structure of the genetic material, each plane is 0.54 μm apart. **D** Imaris spot analysis and annotation of signal positions from Z-stack acquisition. **E** Schematic representation of our interpretation that each signal is a pair of closely associated centromeres from a pair of sister chromatids. As a simplification, all chromosomes have been drawn at an equal length although they actually vary in *O. dioica*.

## Discussion

Despite the variation in signal counts across different image acquisitions settings, a haploid chromosomal count of three provides the most parsimonious explanation of the collected data and agrees with previously published assemblies (Denoeud *et al.*, 2010).

Oocyte staining with rat anti-H3S28P and a conjugated secondary fluorophore gave rise to a compact area in which signals appear to stack on top of one another (Figure 2A). Previously, DNA stains at this stage have been interpreted as a structure resembling the Greek character ∏ (Ganot *et al.* 2007a), representing condensed chromosomes seen in mature oocytes arrested in meiosis I. Our data does not include DNA stains and therefore our illustration (Figure 2E) should not be interpreted as precluding the previously reported ∏-structure.

Currently, the sequence of the centromeres is not known, although chromatin immunoprecipitation with a H3S28P antibody followed by long-read sequencing might be able to provide this information. However, our whole embryo staining data and the previous literature (Table 1) show that non-centromeric signal present outside metaphase stages may introduce noise. Thus, alternative targets such as other centromeric histone 3 variants (Moosmann *et al.*, 2011) might be preferable. Availability of centromeric sequences would open the possibility of confirming our results with fluorescence *in situ* hybridization.

In summary, we conclude that the Okinawan *Oikopleura dioica* genome consists of three pairs of chromosomes in diploid cells. We believe that the images may be useful for examining cell cycle specific changes to chromosome structure and encourage the reuse and reanalysis of our data located in the EBI BioImage Archive (Ellenberg *et al.*, 2018).

## Data Availability

Image acquisitions: Image data are available in the BioImage Archive (https://www.ebi.ac.uk/biostudies/preview/studies/S-BIAD21) under accession number S-BIAD21 and available for use under the CC01.0 Public domain dedication.

## Contributions

Andrew W. Liu* – Investigation, formal analysis, curation, writing (original draft)

Yongkai Tan* – Conceptualization, investigation, methodology, formal analysis, visualization

Aki Masunaga – Visualization, writing (review/editing)

Charles Plessy – Conceptualization, writing (review/editing) & project administration

Nicholas M. Luscombe – Conceptualization, funding acquisition, supervision, writing (review/editing

*Co-first authors

## Acknowledgements

We thank Drs. Daniel Chourrout, Hiroki Nishida & Eiichi Shoguchi for discussions and suggestions regarding the subject matter. Additionally, great appreciation is given the staff (Drs. Toshiaki Mochizuki, Shinya Komoro & Paolo Barzaghi) in the Imaging Section of the Research Support Division at OIST for providing technical assistance. Finally, we are grateful for Dr. Michael Mansfield and Charlotte West comments on the manuscript’s draft and appearance.

## References

Campsteijn C., Øvrebø J.I., Karlsen B. O. and Thompson E.M. (2012) Expansion of Cyclin D and CDK1 Paralogs in *Oikopleura dioica*, a Chordate Employing Diverse Cell Cycle Variants. Mol. Biol. Evol., 29(2):487–502. https://doi.org/10.1093/molbev/msr136

Colombera D. & Fenaux R. (1973) Chromosome form and number in the larvacea. Boll. Zoll., 40:347–353. https://doi.org/10.1080/11250007309429248

Denoeud F., Henriet S., Mungpakdee S., Aury J.M., Da Silva C., Brinkmann H., Mikhaleva J., Olse L.C., Jubin C., Cañestro C., Bouquet J.M., Danks G., Poulain J., Campsteijn C., Adamski M., Cross I., Yadetie F., Muffato M., Louis A., Butcher S., Tsagkogeorga G., Konrad A., Singh S., Jensen M.F., Cong E.H., Eikeseth-Otteraa H., Noel B., Antourard V., Procel B.M., Kachouri-Lafond R., Nishino A., Ugolini M., Chourrout P., Nishida H., Aasland R., Huzurbazar S., Westhof E., Lehrach H., Reinhardt R., Weissenbach J., Roy S.W., Delsuc F., Artiguenave F., Postlethwait J.H., Manak J.R., Thompson E.M., Jaillon O., Du Pasquier L., Boudinot P., Liberles D.A.,Volff J.N., Philippe H., Lenhard B., Crollius H.R., Wincker P.,Chourrout D. (2010) Plasticity of animal genome architecture unmasked by rapid evolution of a pelagic tunicate. Science, 330(6009):1381–1385. https://doi.org/10.1126/science.1194167

Ellenberg J., Swedlow J.R., Barlow M., Cook C.E., Sarkans U., Patwardhan A., Brazma A., Birney E. A call for public archives for biological image data. Nat Methods., 2018 Nov;15(11) 849–854. https://doi.org/10.1038/s41592-018-0195-8

Feng H. and Thompson E.M. (2018) Specialization of CDK1 and cyclin B paralog functions in a coenocystic mode of oogenic meiosis, Cell Cycle, 17(12):1425–1444. https://doi.org/10.1080/15384101.2018.1486167

Feng H., Raasholm M., Moosmann A., Campsteijn C., Thompson E.M. (2019) Switching of INCENP paralogs controls transitions in mitotic chromosomal passenger complex functions. Cell Cycle 18(17):2006–2025. https://doi.org/10.1080/15384101.2019.1634954

Ganot P., Thompson E.M. (2002) Patterning through differential endoreduplication in epithelial organogenesis of the chordate, *Oikopleura dioica*. Dev. Biol., 252(1):59–71. https://doi.org/10.1006/dbio.2002.0834

Ganot P., Bouquet J.M. and Thompson E.M. (2006), Comparative organization of follicle, accessory cells and spawning anlagen in dynamic semelparous clutch manipulators, the urochordate *Oikopleuridae*. Biol. Cell, 98: 389–401. https://doi.org/10.1042/BC20060005

Ganot P., Kallesøe T., and Thompson E.M. (2007a) The cytoskeleton organizes germ nuclei with divergent fates and asynchronous cycles in a common cytoplasm during oogenesis in the chordate *Oikopleura*. Dev. Biol., 302(2):577–590. https://doi.org/10.1016/j.ydbio.2006.10.022

Ganot P., Bouquet J.M., Kallesøe T. and Thompson E.M. (2007b) The *Oikopleura* coenocyst, a unique chordate germ cell permitting rapid, extensive modulation of oocyte production. Dev. Biol., 302(2):591–600. https://doi.org/10.1016/j.ydbio.2006.10.021

Ganot P., Schulmeister A., Thompson E.M. (2008) Oocyte selection is concurrent with meiosis resumption in the coenocystic oogenesis of *Oikopleura*. Dev. Biol., 324(2):266–276. https://doi.org/10.1016/j.ydbio.2008.09.016

Giemsa G. (1902) Färbemethoden für malariaparasiten, Zentralbl. Bakteriol.

Giemsa G. (1904) Eine Vereinfachung und Vervollkommnung meiner Methylenblau-Eosin-Färbemethode zur Erzielung der Romanowsky-Nocht’schen Chromatinfärbung, Centralblatt für Bakteriologie.

Hsu T.C., Benirschke K. (1967) An atlas of mammalian chromosomes. Vol 10, Springer Verlag

Kawajiri A., Yasui Y., Goto H., Tatsuka M., Takahashi M., Nagata K-I. and Inagaki M. (2003) Functional Significance of the Specific Sites Phosphorylated in Desmin at Cleavage Furrow: Aurora-B May Phosphorylate and Regulate Type III Intermediate Filaments during Cytokinesis Coordinatedly with Rho-kinase. Mol. Biol. Cell, 14(4):1489–1500. doi: https://doi.org/10.1091/mbc.e02-09-0612

Kurihara D., Matsunaga S., Kawabe A., Fujimoto S., Noda M., Uchiyama S., Fukui, K. (2006) Aurora kinase is required for chromosome segregation in tobacco BY-2 cells. The Plant Journal, 48: 572–580. https://doi.org/10.1111/j.1365-313X.2006.02893.x

Körner W.F. (1952) Untersuchungen über die gehäusebildung bei appendicularien (*Oikopleura dioica* FOL). Z. Morph. u. Ökol. Tiere, 41(1):1–53. https://doi.org/10.1007/BF00407623

Masunaga A., Liu A.W., Tan Y., Scott A., Luscombe N.M. (2020) Streamlined Sampling and Cultivation of the Pelagic Cosmopolitan Larvacean, *Oikopleura dioica*. J. Vis. Exp, e61279. https://doi.org/10.3791/61279

Moosmann A., Campsteijn C., Jansen P.W., Nasrallah C., Raasholm M., Stunnenberg H.G., Thompson E.M. (2011) Histone variant innovation in a rapidly evolving chordate lineage. BMC Evol. Biol., 11(208). https://doi.org/10.1186/1471-2148-11-208

Olsen L.C., Kourtesis I., Busengdal H., Jensen M.F., Hausen H., Chourrout D. (2018) Evidence for a centrosome-attracting body like structure in germ-soma segregation during early development, in the urochordate *Oikopleura dioica*. BMC Dev. Biol., 18(4). https://doi.org/10.1186/s12861-018-0165-5

Øvrebø J.I, Campsteijn C., Kourtesis I., Hausen H., Raasholm M., Thompson E.M. (2015) Functional specialization of chordate CDK1 paralogs during oogenic meiosis. Cell Cycle, 14(6):880–93. https://doi.org/10.1080/15384101.2015.1006000

Schulmeister A., Schmid M. and Thompson E.M. (2007) Phosphorylation of the histone H3.3 variant in mitosis and meiosis of the urochordate *Oikopleura dioica*. Chromosome Res., 15(189). https://doi.org/10.1007/s10577-006-1112-z

Shoguchi E., Kawashima T., Nishida-Umehara C., Matsuda Y. and Satoh N. (2005) Molecular Cytogenetic Characterization of Ciona intestinalis Chromosomes Zool. Sci., 22(5):511–516. https://doi.org/10.2108/zsj.22.511

Spada F., Vincent M. & Thompson E.M. (2005) Plasticity of histone modifications across the invertebrate to vertebrate transition: Histone H3 lysine 4 trimethylation in heterochromatin Chromosome Res., 13(1):57–72. https://doi.org/10.1007/s10577-005-6845-6

Tjio J., Levan A. (1950) Quadruple Structure of the Centromere. Nature 165, 368 https://doi.org/10.1038/165368a0

